# Differences in effective ploidy as drivers of genome-wide endosperm expression asymmetries and seed failure in wild tomato hybrids

**DOI:** 10.1101/459925

**Authors:** Morgane Roth, Ana M. Florez-Rueda, Thomas Städler

## Abstract

Endosperm misdevelopment leading to hybrid seed failure is a common cause of postzygotic isolation in angiosperms and is observed in both interploidy and homoploid crosses between closely related lineages. Moreover, parental dosage is critical for successful endosperm and seed development, typically requiring a ratio of two maternal to one paternal genome(s) in within-species crosses. The recently revived concept of ‘effective ploidy’ can largely explain the outcome of experimental crosses that (partly) ameliorate hybrid seed failure by manipulating the actual ploidy in one of the parents. However, genome-wide expression perturbations concomitant with levels of hybrid seed failure have yet to be reported. The tomato clade (*Solanum* section *Lycopersicon*), encompassing closely related diploids with partial-to-complete hybrid seed failure and diverse mating systems, provides outstanding opportunities to study these issues. Here we compared replicated endosperm transcriptomes from six crosses within and among three wild tomato lineages. Strikingly, both strongly inviable hybrid crosses displayed conspicuous, asymmetric expression perturbations with strong signatures of cross direction. In particular, *Solanum peruvianum*, the species inferred to have evolved higher effective ploidy than the other two, drove hybrid expression landscapes in both maternal and paternal roles. This global expression divergence was mirrored in functionally important gene families such as transcription factors and E3 ubiquitin ligases, and revealed differences in cell-cycle tuning between lineages that match phenotypic differences in developing endosperm and mature seed size between reciprocal crosses. Our work initiates the exploration of links between parental conflict, genomic imprinting, expression dosage and hybrid seed failure in flowering plants.

## Introduction

Hybrid seed failure (HSF) is a common phenotype mediating early-acting postzygotic reproductive isolation in flowering plants. HSF does not necessarily result from F1 embryo defects as embryos may be rescued from developing seeds and grown to become fertile plants (Sharma *et al.* 1996). Such observations have been widely interpreted as evidence for hybrid endosperms’ compromised ability to correctly nourish the embryo (Lester and Kang 1998; Sekine *et al.* 2013; Rebernig *et al.* 2015). As the products of double fertilization, embryo and endosperm are genetically closely related, yet these fertilization products are strongly dissimilar, concomitant with their different genome composition (embryo diploid, 1m:1p; endosperm triploid, 2m:1p) and methylation profiles (Gehring *et al.* 2009). This original ‘brotherhood’ between endosperm and embryo evolved over long periods of coevolutionary history, which might have contributed to the success of flowering plants (Baroux *et al.* 2002).

The frequent occurrence of HSF in interploidy crosses has been interpreted to be a consequence of endosperm sensitivity to parental dosage, establishing a reproductive barrier termed the ‘triploid block’ (Köhler *et al.* 2010; Stoute *et al.* 2012). A well-known feature of interploid seed failure are the typically contrasting phenotypes of reciprocal developing and/or mature hybrid seeds (Cooper and Brink 1945; Valentine and Woodell 1963; Scott *et al.* 1998; Leblanc *et al.* 2002). Specifically, these asymmetric phenotypes comprise smaller seeds when the ovule parent is of higher ploidy (‘maternal-excess phenotype’) and larger seeds when the pollen parent is of higher ploidy (‘paternal-excess phenotype’; Haig and Westoby 1991). As endosperm size—which largely determines mature seed size—is affected in corresponding directions in such reciprocal interploidy crosses, parental-excess phenotypes have been regarded as a direct consequence of asymmetric parental dosage in their endosperms (Scott *et al.* 1998; Sabelli and Larkins 2009; Stoute *et al.* 2012).

Importantly, such inferred dosage sensitivity is also suspected to play a role in the developmental trajectory and (often) abortion of homoploid hybrid seeds with similar symptoms of parental excess (Josefsson *et al.* 2006; Rebernig *et al.* 2015; Oneal *et al.* 2016; Lafon-Placette *et al.* 2017, 2018). Phenotypic asymmetries between seeds from reciprocal homoploid crosses indicate that incompatibilities expressed in hybrid endosperm encompass parental effects. These phenomena might be caused by differences in so-called ‘effective ploidy, a compound property thought to determine dosage requirements for specific genes in a given lineage (reviewed in Lafon-Placette and Köhler 2016), and in classical work on tuber-bearing *Solanum* species proposed as ‘endosperm balance number’ (EBN; Johnston *et al*. 1980; Ortiz and Ehlenfeld 1992). In crosses between homoploid species with different effective ploidy, the species with higher effective ploidy would mimic the lineage with higher actual (karyotypic) ploidy in an interploidy cross.

From an evolutionary point of view, variation in effective ploidy or ‘genetic strength’ is regarded as a potential consequence of divergence between species in levels of parental conflict. According to this line of thinking, maternal interests ought to restrict seed growth to allocate resources equally among all seeds (from potentially different fathers). In contrast, paternal interests ought to promote growth only for their own sires in the face of other potential fathers, setting up competition for resource allocation between seeds from the same mother (Haig and Westoby 1991; Brandvain and Haig 2005). Under this scenario, the smaller seeds observed in maternal-excess crosses could be a manifestation of growth restrictions of maternal origin, while the larger seeds in paternal-excess crosses might reveal weakened maternal control over resource allocation, thus leading to paternally-driven overgrowth.

Thus far, dissimilar seed phenotypes have been revealed in interploidy crosses, yet without addressing variability in parental conflict strength between lineages. Indeed, while interploidy crosses can reveal parental effects, parental conflict is not expected to depend on ploidy level *per se*. Arguably, studies on homoploid interspecific hybrids are more suitable to investigate whether parental conflict strength has evolved in response to differences in mating system, long-term demographic history, and/or other evolutionary forces. Relevant studies have recently been performed in two Brassicaceae genera, *Arabidopsis* and *Capsella*, where it was shown that the parent with the outcrossing breeding system (*A. lyrata*, *A. arenosa* and *C. grandiflora*, respectively) drives seed phenotypes of maternal- and paternal-excess (Josefsson *et al.* 2006; Rebernig *et al.* 2015; Lafon-Placette *et al.* 2017, 2018); experimentally increasing the ploidy of the inbreeding species partly restored seed viability (Josefsson *et al.* 2006; Lafon-Placette *et al.* 2017). Beyond these phenotypic evidences, divergence in dosage between parental species of flowering plants and its consequences for genome-wide expression modulation appear to not have previously been quantified.

To date, genome-wide studies on endosperm gene expression have mainly focused on characterizing genomic imprinting, *i.e.* the parent-of-origin–dependent expression of genes. A trend for elevated expression of imprinted genes in species with higher parental conflict was found between closely related species, but it is currently not known whether this might contribute to incidences of HSF (Klosinska *et al.* 2016; Roth *et al*. 2018b). Of note, genomic imprinting is extensively perturbed in failing wild tomato hybrid endosperm (Florez-Rueda *et al*. 2016), but it is unclear whether mis-imprinting *per se* or total expression-level changes of functionally important genes (plausibly including imprinted genes) underpin hybrid seed abortion. We may hypothesize that parental imbalances caused by divergent effective ploidies in homoploid crosses affect global expression levels and dosage-sensitive processes such as genomic imprinting. Moreover, we expect such parental imbalances to be reflected by opposite patterns of expression change in the reciprocal crosses.

Wild tomatoes (*Solanum* section *Lycopersicon*) provide a well-suited plant system to study developmental and evolutionary questions on HSF (Florez-Rueda *et al*. 2016; Roth *et al*. 2018a). We have recently shown that crosses between *S. arcanum* var. marañón (A), *S. chilense* (C) and *S. peruvianum* (P) result in different degrees of endosperm disruption leading to partial or complete seed inviability (Roth *et al*. 2018a). In particular, crosses between A and C yield variable proportions of viable and inviable seeds (here categorized as ‘weak-HSF’) whereas crosses between P and either A or C result in near-complete seed failure (termed ‘strong-HSF’; Figure S1). Moreover, marked phenotypic asymmetries are characteristic of seeds from reciprocal crosses of the strong-HSF category, where endosperms fathered by species P (*i.e*. from crosses AP and CP) correspond to paternal-excess phenotypes and endosperms of P maternal plants (*i.e*. from crosses PA and PC) correspond to maternal-excess phenotypes (Florez-Rueda 2014; Roth *et al.* 2018a). We thus hypothesized that lineage P experienced higher levels of parental conflict that led to its increased effective ploidy compared to C and A during their evolutionary divergence.

The present study seeks to (i) quantify molecular perturbations of gene expression levels in (partly or entirely) failing wild tomato endosperm, (ii) assess the prediction of genome-wide asymmetries in patterns of endosperm expression levels between reciprocal strong-HSF crosses, (iii) identify candidate genes/gene families with potentially important roles in expression perturbation, and (iv) discuss the role of parental conflict-driven differences in effective ploidy in causing or contributing to hybrid seed failure.

## Materials and Methods

### Plant material and crosses

Seeds were obtained from the Tomato Genetics Resource Center (TGRC, University of California, Davis, USA; https://tgrc.ucdavis.edu). We crossed three genotypes (one per species) in a full diallele design with all reciprocal crosses producing seed phenotypes typical for weak or strong seed inviability, respectively (Roth *et al.* 2018a; Figure S1). Genotypes were chosen from population LA2185 (Amazonas, Peru) for *S. arcanum* var. marañón (A), population LA4329 (Antofagasta, Chile) for *S. chilense* (C) and population LA2744 (Arica and Parinacota, Chile) for *S. peruvianum* (P) to be used in hybrid crosses (Figure S2). In addition, we chose three genotypes from additional populations of each species in order to perform intraspecific reciprocal crosses (referred to as ‘controls’; Figure S2). The corresponding populations are LA1626 (Ancash, Peru) for A, LA2748 (Tarapaca, Chile) for C and LA2964 (Tacna, Peru) for P. The latter three populations were not used in hybrid crosses. As detailed in Roth *et al*. (2018a), all crosses produced normal quantities of seeds per fruit. Plants were grown from seed in an insect-free greenhouse at ETHZ (Lindau-Eschikon, canton Zurich, Switzerland). They were regularly repotted in 5-l pots using fresh soil (Ricoter Substrate 214, Ricoter Erdaufbereitung AG, Aarberg, Switzerland) and fertilizing granules (Gartensegen, Hauert HBG Dünger AG, Grossaffoltern, Switzerland). Additional liquid fertilizer was applied once or twice per month depending on the season (Wuxal^®^ NPK solution, Aglukon Spezialdünger GmbH and Co. KG, Düsseldorf, Germany). Plants were watered two to four times per week.

Well before the onset of the experiments, cuttings yielded multiple ramets per genotype, from which we chose three to serve as biological replicates. All clones were maintained in a climate chamber for the duration of the whole experiment (12 h light at 18 Klux and 50% relative humidity, 12 h darkness at 0 Klux with 60% relative humidity). Reciprocal crosses were named with the two initial letters of parental lineages within brackets (all reciprocal crosses are: [AC], [AP], [CP], [AA], [CC], [PP]), and individual crosses designated by the initial letters of parental lineages without brackets, indicating the cross direction ‘mother × father’: **AA1**, LA2185A × LA1626B; **AA2**, LA1626B × LA2185A; **CC1**, LA4329B × LA2748B; **CC2**, LA2748B × LA4329B; **PP1**, LA2744B × LA2964A; **PP2**, LA2964A × LA2744B; **AC**, LA2185A × LA4329B; **CA**, LA4329B × LA2185A; **AP**, LA2185A × LA2744B; **PA**, LA2744B × LA2185A; **CP**, LA4329B × LA2744B; **PC**, LA2744B x LA4329B. This implies that AC, AP, and AA1 share the same mother, that CA, CP, and CC1 share the same mother, and that PA, PC, and PP1 share the same mother. Each cross was performed three times using clonal replicates of each genotype. Fruits were sampled 12 days after manual pollinations (12 DAP)—corresponding to the early globular embryo stage—embedded, and endosperms were sampled from fruit cryosections via laser-assisted microdissection. Methods for endosperm sampling, RNA extraction, library preparation and sequencing are detailed in our previous study (Roth *et al*. 2018b).

### Alignment and counting methods

Short read alignment was done as previously described (Roth *et al*. 2018b). Briefly, RNA-Seq quality assessment of all samples was performed with the FastQC program (http://bioinformatics.babraham.ac.uk/projects/fastqc/). Adapters were removed with cutadapt (Martin 2011). Trimming and quality filtering were done with the Perl script trimmingreads.pl from the NGSQC Toolkit version 2.3 (Patel and Jain 2012). Read mapping was performed with TopHat version 2.1.0 (Trapnell *et al.* 2009) against the SL2.50 reference genome of the cultivated tomato var. Heinz (The Tomato Genome Consortium 2012) with the corresponding annotation ITAG2.4 (International Tomato Annotation Consortium; https://solgenomics.net/). Mapping quality check was done with Qualimap version 2.2 (Okonechnikov *et al.* 2016) and RSeQC (Wang *et al.* 2012). Total reads per gene were counted from bam files with HTseq (Anders *et al.* 2015) using the gff ITAG2.4 annotation file (The Tomato Genome Consortium 2012). Only reads with mapping quality above 20 were retained.

### Statistical analyses

Differential gene expression analysis (DGE) was performed with the R package EdgeR (Robinson *et al.* 2010; R Development Core Team 2014). Only genes with at least one read count per million in at least two of the 36 libraries were kept, resulting in a set of 22,006 genes. We used Multidimensional Scaling (MDS) plots to assess variation between biological replicates, using the function plotMDS in EdgeR and target groups ‘species’ for intraspecific crosses and ‘cross type’ for the whole dataset. A negative binomial model was fitted to each gene using individual crosses as factors, estimating trended dispersions (variance parameters). Differentially Expressed Genes (DEGs) were identified in the selected pairwise comparisons using different contrasts with a generalized linear model likelihood ratio test (*P*-value correction with the Benjamini–Hochberg method for a false discovery rate [FDR] of 5%).

In each comparison, we used specific contrasts to compare two classes of crosses according to different criteria: (i) their seed phenotype (*e.g.* in the ‘strong–intra’ comparison, strongly abortive crosses were compared to intraspecific crosses by pooling all replicates of all strong-HSF crosses together (*i.e*. AP, PA, CP, and PC) and comparing them to all replicates of all intraspecific crosses pooled together (*i.e*. AA1, AA2, CC1, CC2, PP1, and PP2)); (ii) hybrids compared to their respective intraspecific cross sharing the same mother (*e.g*. in the ‘PA-PP1’ comparison, all replicates of the PA cross were compared to all replicates of the PP1 cross); (iii) cross direction by comparing reciprocal crosses (*e.g*. in the ‘PA-AP’ comparison, all replicates of the PA cross were compared to all replicates of the AP cross); and (iv) the species in intraspecific crosses (*e.g*. in the ‘[AA]-[CC]’ comparison, we compared all replicates of AA1 and AA2 to all replicates of CC1 and CC2); in total we report 18 different contrasts (Table S1, sheet ‘Contrasts’). Count data used for creating heat maps were obtained from normalized counts per million, averaged across replicates for each cross. Heat maps were plotted with the R package ‘gplots’ using hierarchical clustering (R Development Core Team 2014; Warnes *et al.* 2016). The R package ‘topGO’ (Alexa and Rahnenführer 2016) was used to identify enriched Gene Ontology (GO) terms from ITAG 2.4 downloaded from Plant Ensembl Biomart datamining platform (Kinsella *et al.* 2011), using as gene universe the set of 22,006 endosperm-expressed genes. Venn diagrams were obtained with the R package ‘venneuler’ (Wilkinson 2011).

### Data availability

Raw sequence data for the RNA-sequencing dataset used in this study are available from the Sequence Read Archive (https://trace.ncbi.nlm.nih.gov/Traces/sra/) with the accession numbers PRJNA427095 (18 hybrid endosperm libraries), SRP132466 (18 within-species endosperm libraries and five parental plants; Roth *et al*. 2018b), and SRX1850236 (parent LA4329B; Florez-Rueda *et al*. 2016). Supplemental Material, Figure S1 details the distribution of seed viability in all crosses used in this study, which are a subset of a larger phenotypic study of (hybrid) seed viability (Roth *et al*. 2018a). Figure S2 is a diagram of the crossing design representing the six reciprocal crosses used for our endosperm RNA-Seq experiment. Table S1 is as large Excel table containing four data sheets: ‘Contrasts’ contains the list of 18 comparisons with their corresponding contrasts used in this study, ‘DEGs’ summarizes differential gene expression (DGE) for each of them, ‘GO_enrichment’ summarizes GO-term enrichments for differentially expressed genes (DEGs) in selected categories, and ‘DGE_imprinted_genes’ lists the status of candidate imprinted genes and their differential expression in all tested contrasts.

## Results and Discussion

### Molecular responses to hybridization correspond to hybrid seed failure severity

Seeds with similar phenotypes are likely to have similar expression patterns in the endosperm and low proportions of DEGs between them. In turn, we hypothesized that the magnitude of gene expression differences between two crosses would broadly match their developmental trajectories (along the gradient intraspecific – weak HSF – strong HSF). We assessed this hypothesis with a suite of DGE analyses. After filtering our dataset for lowly expressed genes across the 36 libraries, 22,006 genes remained for DGE analysis, indicating that 63.4% of the ITAG2.4-annotated genes were jointly expressed in the 12-DAP endosperm of our various cross types. The multidimensional scaling (MDS) plot using expression data from only the intraspecific crosses revealed that samples broadly group by species and cross direction (Figure 1A). In particular, differences in the overall gene expression landscape between [CC] and [PP] endosperms appear to be fewer than between [CC] and [AA] or [PP] and [AA] endosperms: 817 DEGs were found between [PP] and [CC], 1,226 DEGs between [CC] and [AA], and 1,184 DEGs between [PP] and [AA]. This apparent genome-wide expression divergence broadly reflects the differences in divergence time between A, C and P (Städler *et al.* 2008; Beddows *et al.* 2017). A positive correlation between genomic and expression divergence is expected from theory (Nuzhdin *et al.* 2004; Renaut *et al.* 2012). However, while our results support this notion, the correlation between expression and sequence divergence appears to be either positive (Nuzhdin *et al.* 2004; Khaitovich *et al*. 2005; Renaut *et al.* 2012) or non-significant (Jeukens *et al.* 2010; Wolf *et al.* 2010; Moyers and Rieseberg 2013) in previous empirical studies.

**Figure 1.**
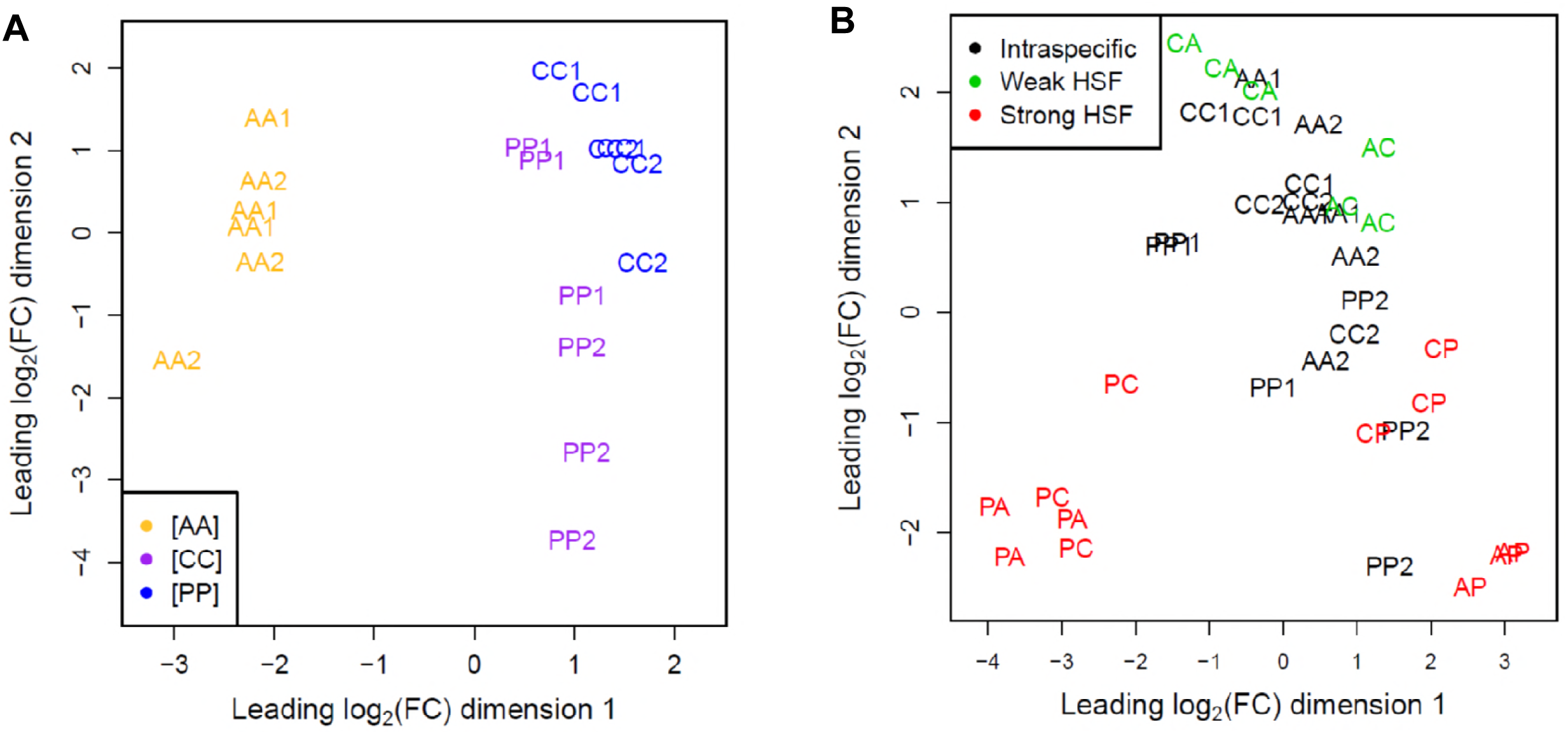
Multidimensional scaling plot representing the distances between endosperm samples (*i.e.* sequencing libraries) based on the joint expression levels of 22,006 genes. (A) All 18 samples representing intraspecific, reciprocal crosses [AA], [CC] and [PP]. (B) All 36 endosperm samples (*i.e.* intraspecific as well as hybrid) analyzed jointly. HSF, hybrid seed failure; log_2_(FC), log_2_-fold change. A, *S. arcanum* var. marañón; C, *S. chilense*; P, *S. peruvianum*. **AA1**, LA2185A × LA1626B; **AA2**, LA1626B × LA2185A; **CC1**, LA4329B × LA2748B; **CC2**, LA2748B × LA4329B; **PP1**, LA2744B × LA2964A; **PP2**, LA2964A × LA2744B; **AC**, LA2185A × LA4329B; **CA**, LA4329B × LA2185A; **AP**, LA2185A × LA2744B; **PA**, LA2744B × LA2185A; **CP**, LA4329B × LA2744B; **PC**, LA2744B × LA4329B. Cross specifications are identical in all other display items.

The global expression landscape represented by the joint analysis of all 36 samples revealed several expression profiles corresponding to different seed phenotypes (Figure 1B); the y-axis mainly separates intraspecific and weak-HSF crosses [AA], [CC], [PP] and [AC] from strong-HSF crosses [AP] and [CP]. We quantified these expression changes and found that many more genes are differentially expressed between strong-HSF ([AP] and [CP]) and intraspecific endosperms ([AA], [CC] and [PP]) than between weak-HSF ([AC]) and intraspecific endosperms (3,026 vs. 682 DEGs; Figure 2A; Table S1, sheet ‘DEGs’). Interestingly, 85.5% of DEGs between strong-HSF and intraspecific endosperms overlap with DEGs between strong-HSF and weak-HSF endosperms (with the same direction of expression changes relative to strong-HSF endosperms; Table S1, sheet ‘DEGs’). Expression differences in hybrid endosperms are likely a product of hybridization *per se* (Hegarty *et al.* 2009; Combes *et al.* 2015; Raza *et al.* 2017), but expression perturbation may also be expected to be stronger when parental species are more genetically diverged (Landry *et al.* 2007; Stelkens and Seehausen 2009; He *et al.* 2010). Because expression differences among intraspecific crosses reflect genetic divergence between lineages (Figure 1A), we might have expected [CP] to exhibit the least-altered expression pattern among all hybrids. To the contrary, [CP] and [AP] hybrid endosperms revealed the most dissimilar expression patterns compared to their parental intraspecific crosses, while [AC] endosperms were close to their parental intraspecific crosses in terms of overall expression landscape (Figures 1B, 2B). Also, interspecific expression differences contributed more to DEGs observed between individual hybrids and their corresponding intraspecific cross (sharing the same mother) in weak-HSF hybrids (52.5–54.4%) than in strong-HSF hybrids (only 19.3–31.9%). This suggests that gene expression divergence between parental species alone cannot explain the extensive expression changes in strongly abortive endosperms (Figure 3); rather, epistatic interactions might rewire gene regulation and generate unique expression landscapes in [AP] and [CP] hybrids which might be responsible for their extreme phenotypes and near-complete inviability (Renaut *et al.* 2009; Dion-Côté *et al.* 2014; Roth *et al*. 2018a).

**Figure 2.**
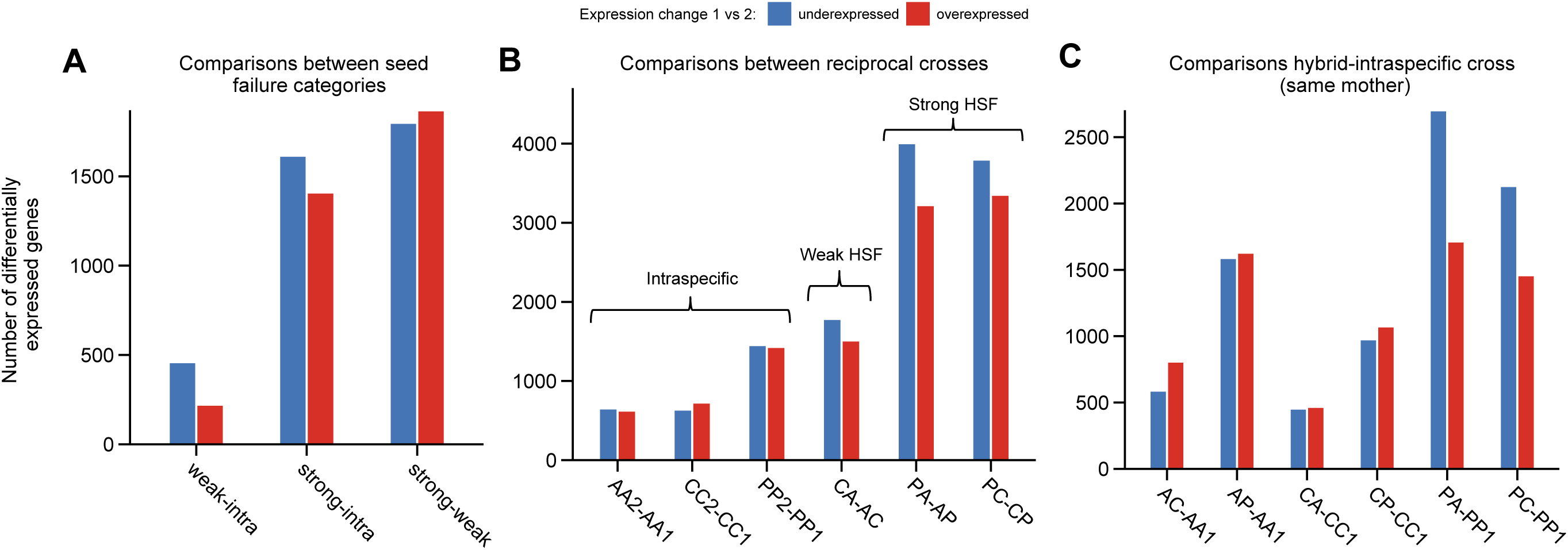
Overview of numbers of differentially expressed genes (DEGs) in different cross comparisons. The direction of expression change refers to the first term as compared to the second term (e.g. genes upregulated in ‘weak-intra’ refers to genes upregulated in ‘weak’ compared to ‘intra’). Blue, underexpressed genes; red, overexpressed genes. (A) DEGs between designated classes of seed failure phenotype (i.e. intraspecific, weak hybrid seed failure (HSF), strong HSF). (B) DEGs between reciprocal crosses. (C) DEGs between hybrid and intraspecific crosses sharing the same mother.

**Figure 3.**
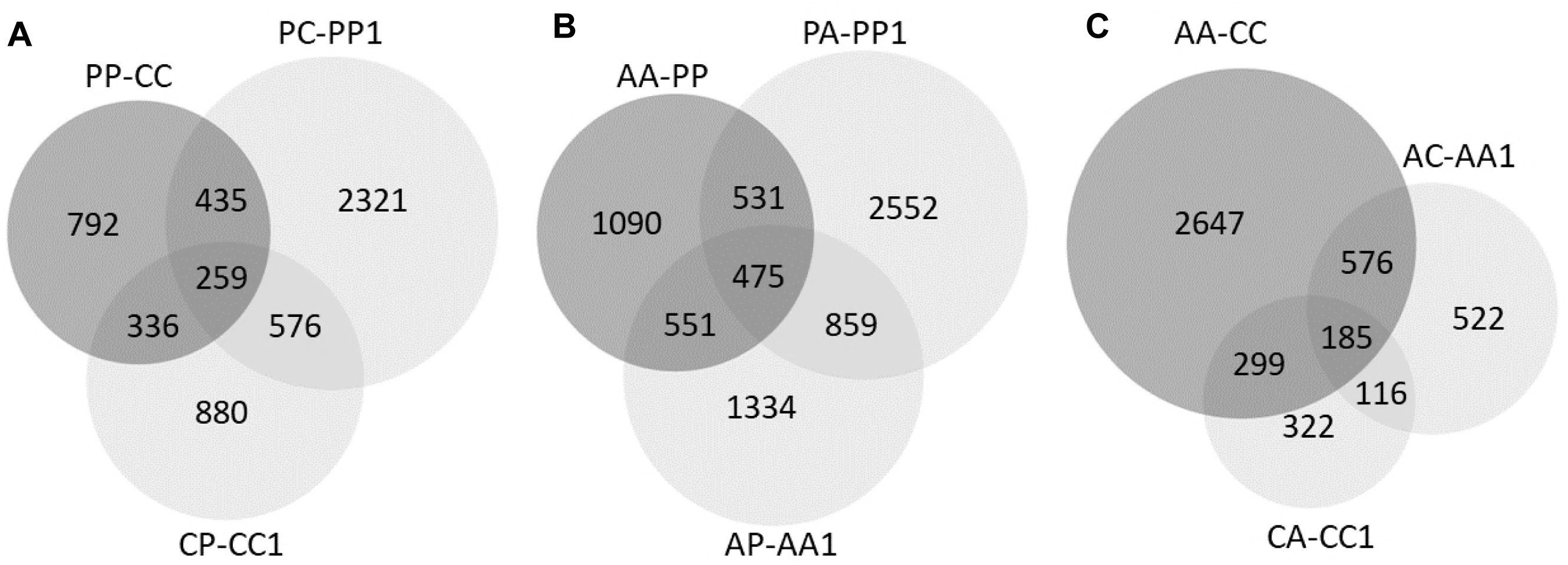
Venn diagrams representing the overlap between DEGs identified in different cross comparisons. (A) Comparisons involving lineages C and P; (B) comparisons involving lineages A and P; (C) comparisons involving lineages A and C.

As a consequence of extensive transcriptomic changes, a wide range of biological functions were affected in strong-HSF endosperms. DEGs between strong-HSF and intraspecific endosperms were enriched for carbohydrate and lipid metabolism, transcription regulation, chromatin conformation, cell cycle, cell structure (cell wall, microtubules), signalling (peptides, hormones, response to stress) and transport (100 GO terms; Table S1, sheet ‘GO_enrichment’). The far fewer DEGs between weak-HSF and intraspecific endosperms were mainly enriched for terms related to carbohydrate and lipid metabolism (34 GO terms; Table S1, sheet ‘GO_enrichment’). Changes related to signalling and cell wall modifications have been reported as potential contributors to HSF in *Arabidopsis* hybrid endosperm (Burkart-Waco *et al.* 2013), but functions relating to global transcriptome changes during endosperm-based HSF remain poorly documented. Interestingly, functions enriched among imprinted genes (*i.e.* those with parent-of-origin–dependent expression) whose expression levels may be critical for seed development, seem to overlap with perturbed functions observed in strong-HSF endosperms. In particular, ten enriched GO terms found among strong-HSF DEGs were in common with GO terms enriched among wild tomato imprinted genes (Roth *et al*. 2018b; Table S1, sheet ‘GO_enrichment’). These GO terms correspond mainly to transcription factor activity, metabolic processes and signalling and are also found enriched among imprinted genes in other species such as *A. thaliana*, rice, maize and sorghum (Gehring *et al.* 2011; Luo *et al.* 2011; Waters *et al.* 2013; Zhang *et al.* 2016). Focusing on 58 conserved imprinted genes previously identified in our three wild tomato species (Roth *et al*. 2018b), we found that 23 were differentially expressed in strong-HSF endosperm vs. only four in weak-HSF endosperm, when compared to intraspecific crosses (Table S1, sheet ‘DGE_imprinted_genes’). While we have previously shown that maternal-to-paternal expression ratios are markedly perturbed in abortive [CP] crosses (Florez-Rueda *et al*. 2016), our results demonstrate that total expression levels of imprinted genes are also affected and could contribute to HSF in strong-HSF crosses.

### Expression asymmetries between reciprocal crosses match parental-excess phenotypes

For the entire transcriptome data set, we found the strongest expression differences between the reciprocals of strong-HSF crosses. Indeed, the expression landscapes of crosses with P as the ovule parent (PA and PC) on the one hand and crosses with P as the pollen parent (AP and CP) on the other hand, are fundamentally dissimilar (x-axis of the MDS plot; Figure 1B). This marked expression divergence corresponds to opposite seed phenotypes, comprising larger seeds in AP and CP crosses (‘paternal excess’-like) and smaller seeds in PA and PC crosses (‘maternal excess’-like; see Introduction).

The DGE analysis revealed that about one third of all endosperm-expressed genes were differentially expressed between both the PA and AP (*n* = 7,227 genes) and the PC and CP crosses (*n* = 7,153 genes; Figure 2C). This indicates profound parental dosage differences between reciprocals which qualitatively inherit the same parental genomes but differ in the dosage from each parent due to the asymmetric 2m:1p endosperm genomic content. In both the [AP] and [CP] reciprocal crosses, more genes were overexpressed when P was in the paternal than when it was in the maternal role (Figure 2C). Moreover, of these two sets of DEGs, 4,477 genes were in common and shared the same direction of expression change in both the ‘PA-AP’ and ‘PC-CP’ comparisons (only 127 genes showed opposite gene expression changes between them). This high proportion of shared gene identity and expression change implies that the strong-HSF endosperms respond in a highly symmetric fashion relative to parent P, indicating that the relative dosage of P (two as ovule parent and one as pollen parent) drove global expression changes between these two reciprocal hybrid crosses. We performed a functional enrichment analysis for the 4,320 GO-annotated DEGs shared in the two comparisons ‘PA-AP’ and ‘PC-CP’ (with the same direction of expression change with respect to the P parent; Table S1, sheet ‘GO_enrichment’). Overall, these DEGs were mainly enriched for expression regulation, chromatin modifications and a large number of biosynthetic and catalytic processes (Table 1; Table S1, sheet ‘GO_enrichment’). Transcription was affected from initiation to RNA maturation (DNA binding, RNA polymerase II, tRNA, snRNA, posttranscriptional regulation of gene expression; Table 1; Table S1, sheet ‘GO_enrichment’). The expression of genes relating to chromatin modifications was also highly divergent between these crosses (helicases, nucleosome, replication initiation, chiasma assembly; Table 1; Table S1, sheet ‘GO_enrichment’).

**Table 1.** Top 10 GO-terms enriched among genes differentially expressed in reciprocal PA-AP and PC-CP hybrid endosperm.

| Direction of change | Ontology category | GO-term ID | GO term description | #Annotated genes | #Observed genes | #Expected genes | Corrected *P*-value |
| --- | --- | --- | --- | --- | --- | --- | --- |
| Overexpressed when P is father (AP and CP crosses) | MF | 4,161 | dimethylallyltranstransferase activity | 20 | 20 | 2 | 1.75E-17 |
|  | MF | 3,677 | DNA binding | 1,358 | 247 | 163 | 3.25E-15 |
|  | MF | 8,234 | cysteine-type peptidase activity | 133 | 49 | 16 | 2.00E-11 |
|  | MF | 46,983 | protein dimerization activity | 354 | 90 | 43 | 1.63E-09 |
|  | BP | 6,334 | nucleosome assembly | 41 | 20 | 5 | 7.75E-08 |
|  | BP | 6,508 | proteolysis | 684 | 122 | 78 | 7.75E-08 |
|  | MF | 8,289 | lipid binding | 125 | 33 | 15 | 1.50E-07 |
|  | MF | 1,104 | RNA polymerase II transcription cofactor activity | 28 | 13 | 3 | 5.58E-05 |
|  | MF | 46,982 | protein heterodimerization activity | 118 | 31 | 14 | 1.14E-04 |
|  | MF | 30,599 | pectinesterase activity | 49 | 17 | 6 | 1.94E-04 |
| Overexpressed when P is mother (PA and PC crosses) | MF | 3,700 | transcription factor activity | 473 | 98 | 42 | 1.95E-14 |
|  | MF | 43,565 | sequence-specific DNA binding | 265 | 62 | 23 | 3.75E-11 |
|  | MF | 8,146 | sulfotransferase activity | 18 | 11 | 2 | 7.17E-07 |
|  | BP | 6,355 | regulation of transcription, DNA-templated | 1,086 | 143 | 98 | 2.05E-06 |
|  | MF | 45,735 | nutrient reservoir activity | 30 | 13 | 3 | 6.63E-06 |
|  | BP | 9,734 | auxin-activated signaling pathway | 48 | 17 | 4 | 4.75E-05 |
|  | MF | 4,722 | protein serine/threonine phosphatase activity | 79 | 20 | 7 | 1.20E-04 |
|  | MF | 8,289 | lipid binding | 125 | 25 | 11 | 2.08E-04 |
|  | MF | 4,674 | protein serine/threonine kinase activity | 413 | 61 | 36 | 4.36E-04 |
|  | BP | 6,468 | protein phosphorylation | 934 | 122 | 84 | 7.67E-04 |

MF, molecular function; BP, biological process.

DEGs between reciprocal crosses can reveal functions preferentially controlled by one parent that are perturbed in hybrid endosperms. For example, transcription and chromatin-related activities were more often—but not exclusively—enriched among genes overexpressed with P as pollen parent (Table S1, sheet ‘GO_enrichment’). Other functions seemed to be more specifically overexpressed when P was the ovule parent, such as energy metabolism (*e.g*. starch and lipids), stress signals, cell-cycle control (protein phosphorylation, protein serine/threonine kinase and auxin-related terms) and cell architecture (cell wall; Table 1; Table S1, sheet ‘GO_enrichment’). Also, important functional categories among candidate imprinted genes such as DNA-binding (Waters *et al.* 2011; Roth *et al.* 2018b) were enriched among DEGs between reciprocal strong-HSF crosses (Table 1; Table S1, sheet ‘GO_enrichment’). We found that a large proportion of wild tomato conserved imprinted genes were differentially expressed between CP and PC (39 out of 58 genes) and between AP and PA (29 out of 58 genes). In particular, maternally expressed genes (MEGs) were mostly overexpressed in maternal-excess endosperms (32/32 differentially expressed MEGs overexpressed in PC-CP and 20/22 MEGs differentially expressed MEGs overexpressed in PA-AP) while paternally expressed genes (PEGs) tended to be overexpressed in paternal-excess endosperms (7/7 differentially expressed PEGs overexpressed in CP-PC and 5/7 differentially expressed PEGs overexpressed in AP-PA). These expression patterns might indicate that parental excess alters specific dosage mechanisms regulating the expression of imprinted genes, which is potentially lethal for the endosperm and thus the developing seed (Lafon-Placette *et al*. 2018).

Because transcription regulation seems to be deeply affected in reciprocal strong-HSF crosses, we scrutinized expression changes of transcription factors (TFs) and found extensive expression changes among WRKY and MADS-Box TFs (Figure 4A, B). In the WRKY-annotated genes, expression was homogeneous between intraspecific and weak-HSF crosses, but markedly different in strong-HSF endosperms. Two sets of genes were respectively over- and underexpressed in all strong-HSF crosses when compared to intraspecific and weak-HSF crosses. Two other sets of genes exhibited different expression levels between reciprocals of [AP] and [CP] (Figure 4A). The WRKY TF family is very diverse and involved in several major developmental processes including seed development (Rushton *et al.* 2010). One WRKY TF, *TRANSPARENT TESTA GLABRA2*, has been reported as a MEG in *A. thaliana* for which accession-specific dosage is essential for seed survival and involved in the control of endosperm cellularization (Dilkes *et al.* 2008). Among MADS-Box TFs, we also observed two subsets of over- and underexpressed genes in paternal-excess endosperms AP and CP compared to all other cross categories (Figure 4B). MADS-Box genes such as *AGAMOUS-LIKE* (AGL) genes are linked to the Polycomb Repressive Complex (PRC) and involved in *A. thaliana* endosperm cellularization during development (Kang *et al.* 2008; Walia *et al.* 2009). The paternal-excess phenotype of *A. thaliana* × *A. arenosa* interspecific seeds has been linked to the overexpression of several AGL genes in the developing endosperm (Walia *et al.* 2009), while downregulation of *AGL62* in *A. thaliana osd1* mutants results in a maternal-excess phenotype (Kradolfer *et al.* 2013).

**Figure 4.**
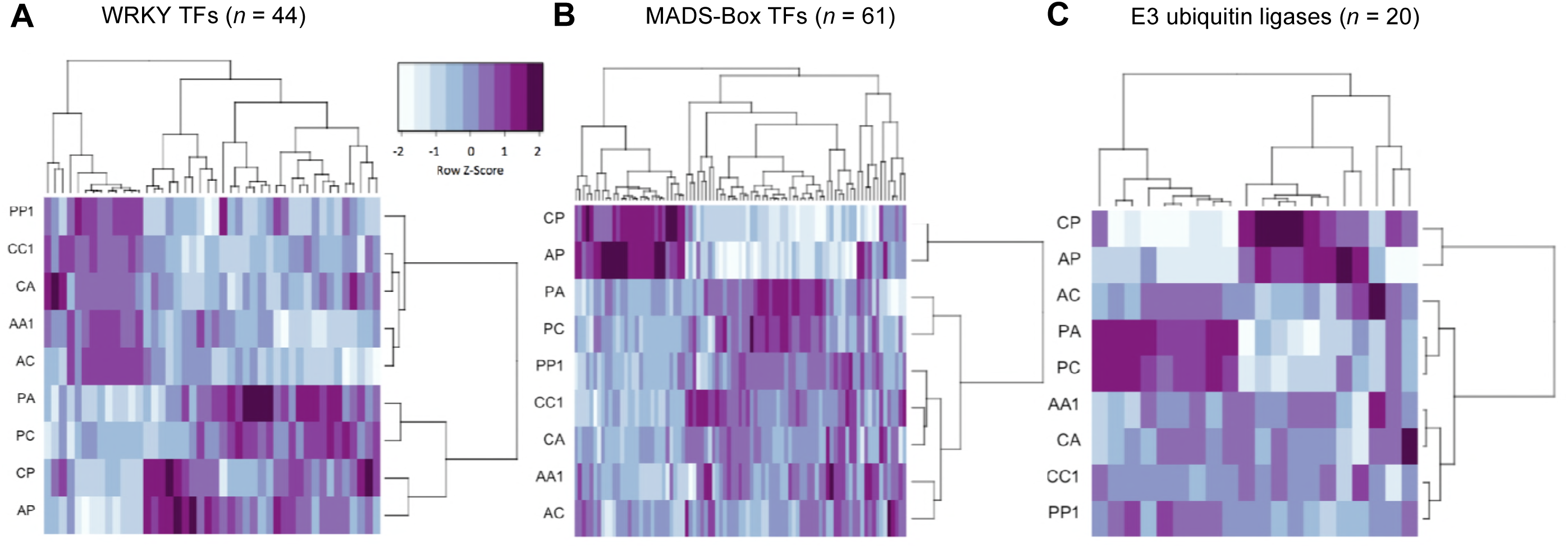
Heat maps representing expression variability for selected gene families among intraspecific and hybrid crosses sharing the same mother. (A) WRKY transcription factors (TFs); (B) MADS-Box TFs; (C) E3 ubiquitin ligases. Color scale according to Z-score (darker colors correspond to stronger expression values); samples and genes ordered by hierarchical clustering.

### Higher effective ploidy of P might underlie phenotypic and transcriptomic asymmetries

The phenotypic asymmetries between reciprocal, inviable hybrid crosses [AP] and [CP] coincide with seed phenotypes typically observed in interploidy crosses (see Introduction). However, P does not have an increased ploidy as all three species studied here are diploid. Although P does not exhibit higher genome-wide expression levels (Roth *et al*. 2018b), our DGE analysis found more genes overexpressed than underexpressed in [PP] endosperm compared to either [AA] or [CC] endosperm, with an overlap of 390 genes overexpressed in [PP] in both comparisons (PP-AA up = 1,471; PP-AA down = 1,176; PP-CC up = 971; PP-CC down = 851; Table 2; Table S1, sheet ‘DEGs’). This indicates that compared to both A and C, lineage P features increased expression in the endosperm that is not observed genome-wide but rather restricted to a subset of genes. Interestingly, among the common set of 390 genes overexpressed in [PP] compared to both [CC] and [AA], a sizable fraction (*n* = 252, 64.6%) comprises genes either overexpressed in both maternal-excess crosses (PA and PC compared to AP and CP, *n* = 129) or overexpressed in both paternal-excess crosses (AP and CP compared to PA and PC, *n* = 123; Table S1, sheet ‘DEGs’). From these sets of genes, genes overexpressed in maternal-excess crosses are mainly enriched for nutrient reservoir activity (*P* = 0.0145) and galactose metabolism (*P* = 0.0037), and genes overexpressed in paternal-excess crosses are enriched for DNA binding (*P* = 3.00e-05), transcription regulation (*P* = 8.00e-05) and biosynthetic process (*P* = 3.55e-05). These enrichments possibly reflect increased maternal influence on resource allocation in maternal-excess endosperms and increased paternal influence on the control of gene expression and growth, respectively.

**Table 2.**
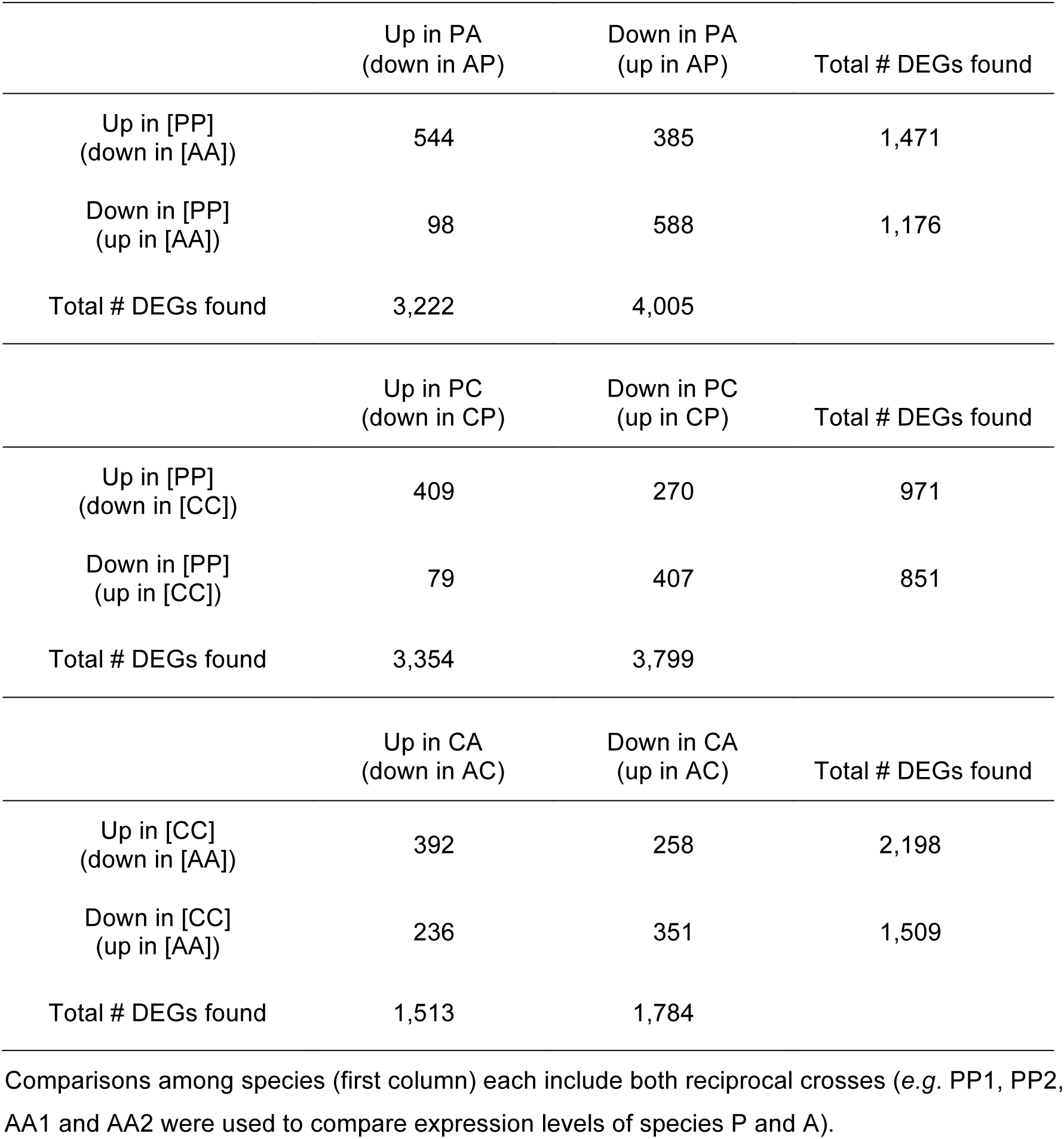
Contingency table of differentially expressed genes (DEGs) in among-species vs. reciprocal hybrid comparisons.

It has recently been shown that imprinted genes in *Capsella*, and especially PEGs, tend to have increased expression in species with the highest effective ploidy (Lafon-Placette *et al*. 2018). In our recent study on wild tomatoes, only a small fraction of candidate imprinted genes were significantly differentially expressed between [AA], [CC] and [PP], and these were exclusively MEGs (Roth *et al*. 2018b; Table S1, sheet ‘DGE_imprinted_genes’). MEGs overexpressed in [PP] were mostly found to be overexpressed in maternal-excess crosses PA and PC (6 of 7 in PA and 4 of 5 in PC; Table S1, sheet ‘DGE_imprinted_genes’). Thus, an increased expression of imprinted genes in P might contribute only marginally to expression asymmetries observed between reciprocals of the strong-HSF crosses. Alternatively, the contribution of imprinted genes might be underestimated because more imprinted genes still remain to be identified due to technical limitations (*e.g.* lack of parental polymorphism for many genes in the crosses used; Roth *et al*. 2018b).

Among the 41 putative AGL genes expressed in wild tomato endosperm, we found that 28 were jointly overexpressed in both paternal-excess crosses (30 of 34 DEGs in CP-PC and 29 of 31 DEGs in AP-PA comparisons). Among them, eight were also overexpressed in [PP] compared to both [CC] and [AA] endosperms. This pattern of expression suggests that these eight AGL genes might be paternally expressed, but their imprinting status could not be assessed due to lack of SNPs between our parental plants (only one AGL gene was polymorphic in P and not imprinted; Roth *et al*. 2018b). Yet, AGL genes are potentially subject to imprinting in the endosperm, as shown by the first-ever identified PEG *PHERES1*, and further AGL genes being maternally or paternally expressed in *Arabidopsis* (Köhler *et al.* 2003; Shirzadi *et al.* 2011; Bai and Settles 2015). Overall, our data indicate increased expression levels in species P for genes known to be critical for seed size and seed viability, such as AGL genes. This might reflect the increased effective ploidy of this species as an evolutionary consequence of higher levels of parental conflict in P compared to both A and C (Lafon-Placette and Köhler 2016; Lafon-Placette *et al*. 2018; Roth *et al.* 2018b).

### Molecular functions underlying parental excess reveal differences in cell-cycle tuning between lineages

GO terms associated with genes differentially expressed between maternal- and paternal-excess crosses (*i.e*. PA and PC versus AP and CP, respectively) indicated contrasting cell cycle regimes. DNA replication and chiasma assembly were enriched among genes overexpressed in paternal-excess endosperms, indicating that AP and CP endosperm cells were probably still dividing at 12 DAP, while proliferation had most likely stopped in the corresponding PA and PC endosperms (Table S1, sheet ‘GO_enrichment’). Our previously published morphological measurements of various [CP] seed compartments between 10 and 13 DAP bolster this inference (Roth *et al.* 2018a). Also, the enrichment in cell cycle control- and cell wall-related terms in DEGs between hybrid endosperms with P in the maternal vs. paternal role (*i.e*. PA and PC versus AP and CP, respectively) is plausibly linked to cell-proliferation differences observed between these endosperms. Related to this, we found striking expression asymmetries in E3 ubiquitin ligases whose protein products are involved in the control of the cell cycle (Inzé and De Veylder 2006; Figure 4C). Among the 20 E3 ubiquitin ligase genes expressed in our data set, eight were overexpressed in maternal-excess compared to paternal-excess endosperms, while only one gene was differentially expressed between the weak-HSF cases CA and AC (Table S1, sheet ‘DEGs’).

The function ‘negative regulation of growth’ was overexpressed in maternal-excess phenotypes, combined with an increased response to auxin (Table S1, sheet ‘GO_enrichment’) which is known to exert negative control of cell division (John *et al.* 1993; Schruff *et al.* 2006; Orozco-Arroyo *et al.* 2015). *A. thaliana arf* mutants bear a non-functional *AUXIN RESPONSE FACTOR 2* (ARF2) and a paternal-excess phenotype with enlarged seeds due to delayed and extended cell divisions in seed tissues (Schruff *et al.* 2006). This indicates that maternal factors control the response to auxin, which is responsible for the control of cell cycle transitions. Interestingly, five ARFs were found to be MEGs in wild tomato endosperm, and three of them were overexpressed in maternal-excess phenotypes (Solyc04g081240.2, Solyc07g043610.2, Solyc11g069500.1; Roth *et al*. 2018b; Table S1, sheet ‘DEGs’). Signals for cell differentiation and response to hormones involved in cell differentiation and seed maturation, such as brassinosteroids and abscisic acid (Orozco-Arroyo *et al.* 2015), were overrepresented among genes overexpressed in the maternal-excess endosperms (Table S1, sheet ‘GO_enrichment’). Compared to intraspecific PP1 endosperm, genes involved in mitotic chromosome condensation and regulation of G2/M transition of the mitotic cell cycle were mainly underexpressed in PA and PC (category ‘down in PA-PP1 & PC-PP1’; Table S1, sheet ‘GO_enrichment’), whereas genes involved in fruit ripening and seed dormancy were overexpressed (category ‘up in PA-PP1 & PC-PP1’; Table S1, sheet ‘GO_enrichment’). These concomitant expression changes probably reflect our histological observations that maternal-excess endosperms stopped dividing and already started to differentiate at the early globular embryo stage (Roth *et al*. 2018a).

We thus suggest that hormone concentrations, regulating the progression through the cell cycle, are mainly under maternal control and perturbed in opposite ways in (PA, PC) versus (AP, CP) endosperms, contributing to maternal- and paternal-excess endosperm morphologies and the corresponding seed size differences. As proposed for interploid maize crosses by Leblanc *et al.* (2002), parental dosage would influence the cell cycle such that (i) rapid mitotic arrest is due to fast G/M transitions in maternal-excess endosperm, and (ii) a longer phase of cell proliferation is due to facilitated re-entry into the S-phase (DNA replication phase) and delayed G/M transitions in paternal-excess endosperm.

Further, some authors have argued that HSF due to dosage imbalance is not a result of perturbed imprinting *per se* but rather a sign that imprinted regulators of cytoplasmic growth rate are misexpressed (von Wangenheim and Peterson 2004; Li and Dickinson 2010). All eukaryotic cells progress through the cell cycle by means of precise control of cyclin concentrations. Although cyclins and their regulation are only partially characterized in plants, it is known that genes encoding cyclins are controlled by growth hormones (Inzé and De Veylder 2006, and references therein). Parental control of hormones and other cell-cycle regulators would support the hypothesis that imprinting evolved to ensure stable production of certain cell components (Hurst and McVean 1998; Weisstein and Spencer 2003). Also, pervasive maternal control over hormone supply could be interpreted as a coadapted control of cell signalling between the endosperm and maternal tissues, thus allowing their synchronized development (Wolf and Hager 2006).

### Parental excess in the endosperm mediates perturbed growth of maternal seed tissues

We previously reported that sporophytic tissues were affected by the hybrid state, notably in CP crosses where both nucellus and seed coat were enlarged compared to [CC] developing seeds (Roth *et al*. 2018a). Based on studies of *A. thaliana arf* mutants, Schruff *et al.* (2006) proposed that impaired auxin regulation in sporophytic tissues altered seed size. As highlighted by our morphological data on some of the same wild tomato hybrid crosses studied here (Roth *et al*. 2018a), seed size and development was impaired in strong-HSF hybrids in a fashion similar to that observed by Schruff *et al*. (2006). However, in our study CP, CA and CC1 seeds inherited the same sporophytic genome (from mother C1) yet exhibited different seed viability levels, suggesting that the perturbation of auxin control is unlikely to originate in sporophytic tissues. Rather, auxin control is most likely first impaired in the endosperms of [AP] and [CP] hybrids and might subsequently mediate hormonal perturbation in sporophytic tissues.

Recent studies in *A. thaliana* have demonstrated that the endosperm-expressed AGL62 mediates the transport of auxin from the endosperm to the integuments and underlies nucellus degradation as well as integument initiation and growth during seed development (Figueiredo *et al*. 2016; Xu *et al.* 2016; Fiume *et al.* 2017). These results strengthen the hypothesis that AGL and auxin deregulation in the endosperm might be tightly linked and indicate that expression perturbation in the endosperm might trigger physiological abnormalities in maternal compartments of wild tomato abortive seeds. Our results thus emphasize the central role of the endosperm as a coordinator of seed development and growth, and they lend support to maternal– offspring coadaptation of gene expression in the seed (Berger *et al.* 2006; Wolf and Hager 2006; Nowack *et al.* 2010).

### Evolutionary implications of differences in effective ploidy for reproductive isolation

We described the transcriptomes of hybrid endosperms obtained by reciprocally crossing three homoploid, closely related wild tomato lineages and found that transcriptomic differences were associated with phenotypic differences between intraspecific, partially viable, and completely inviable hybrid seeds. Our study system included two crosses with reciprocal strong-HSF phenotypes ([AP] and [CP]) which also exhibited similar expression signatures. Thus, the [AP] and [CP] data reflect independently evolved yet similar biological features, suggesting shared molecular and physiological underpinnings of reproductive isolation between closely related lineages.

Moreover, the asymmetric phenotypes and expression landscapes of strongly abortive hybrid seeds indicate that parental conflict has facilitated the establishment of reproductive isolation. More specifically, species P appears to drive HSF at both molecular and phenotypic levels upon hybridization with lineages C or A. We thus propose that P bears an increased effective endosperm dosage, which can be interpreted as a higher effective ploidy (or EBN; Johnston *et al.* 1980; Lafon-Placette and Köhler 2016). Recent empirical data in *Capsella* suggest a positive correlation between levels of parental conflict within lineages and effective ploidy (Rebernig *et al.* 2015; Lafon-Placette *et al.* 2018). As levels of parental conflict should negatively correlate with relatedness between parents, such conflict is expected to decrease with more intense inbreeding (Brandvain and Haig 2005). Although our study included only obligate outcrossers, lineages A, C and P harbor different levels of range-wide nucleotide diversity; specifically, P is the most diverse and A the least diverse lineage (Städler *et al*. 2008; Tellier *et al*. 2011; Labate *et al*. 2014). Range-wide nucleotide diversity should reflect long-term effective population size; all other things being equal, one would expect lower parental conflict between two randomly drawn plants from the least polymorphic (A) compared to the more polymorphic lineages (C and, particularly, P). In summary, we infer the relative effective ploidies between lineages to be P >> C ≥ A.

Lafon-Placette *et al*. (2018) identified higher numbers and expression levels of PEGs in the obligatory outcrosser *Capsella grandiflora* (inferred to have the highest effective ploidy), compared to the highly selfing species *C. rubella* and the more ancient selfer *C. orientalis*. In contrast, our present and previously reported data (Roth *et al*. 2018b) entail that PEGs are expressed at similar levels between A, C and P. We also found no significant differences in the proportion of PEGs between A, C and P (χ^2^ test, *P* > 0.05). This lack of a clear signal regarding the number and expression level of PEGs concomitant with apparent divergence in effective ploidy within our study system can be reconciled due to the presumably closer levels of parental conflict among our wild tomato lineages (with A, C and P all being obligate outcrossers), compared to the *Capsella* system (Lafon-Placette *et al*. 2018).

Hybrid crosses between A and C produced variable proportions of viable seeds, suggesting they have roughly comparable effective ploidies. Despite the occurrence of a few developmental abnormalities, germinating [CA] F1 hybrids proved viable (Roth *et al*. 2018a), indicating that lineages C and A have accrued only few genetic incompatibilities. On the other hand, in the crosses selected for the present study, AC seeds were larger than [AA] seeds and much larger than CA seeds, suggesting a pattern of paternal excess for AC seeds (Roth *et al*. 2018a). Consequently, C manifests signs of higher effective ploidy compared with A, but this dosage difference appears small enough to be overcome by natural (endosperm) robustness to hybridization, at least for a large fraction of seeds. Unfortunately, our dissection protocol does not allow discriminating the endosperm from viable and non-viable seeds at the pre-globular embryo stage, which could be useful to compare the transcriptomes of non-viable and viable hybrids seeds from ‘weak-HSF’ crosses.

Importantly, we did not find significant genome-wide differences in expression levels between [AA], [CC] and [PP] endosperms and relatively few DEGs between them (Figures 1A, 3; Table 2; Table S1, sheet ‘DEGs’). Hence, we propose that the property ‘effective ploidy’ manifests as the stronger expression of a limited number of specific genes controlling dosage-sensitive mechanisms, such as cell-cycle regulation; we provide a number of candidate mechanisms controlling this feature. In particular, expression levels of AGL genes seem to match the inferred genetic-value hierarchy; among the 41 putative AGL genes expressed in our dataset, eight showed expression differences between the intraspecific crosses [AA], [CC] and [PP], such that P > C, P > A, and C > A. All eight genes were overexpressed in the paternal-excess endosperms AP and CP, and four of them were also overexpressed in AC (the hybrid combination exhibiting ‘milder’ paternal excess) such that AP > PA, CP > PC, and AC > CA (Table 3). These eight AGL genes are thus candidates for underpinning different effective ploidies between tomato lineages, and as a consequence they might be decisive for the occurrence and/or severity of HSF. Knocking out single or multiple AGL genes in parental plants or modifying their expression levels in the endosperm, as has been done in *Arabidopsis* (Walia *et al.* 2009; Kradolfer *et al.* 2013), would allow to validate their specific roles (if any) in endosperm development and seed failure in *Solanum*. The expression patterns of AGL genes in intraspecific and reciprocal hybrid comparisons indicate that they might be paternally expressed, but imprinting could not be assessed for these genes.

**Table 3.**
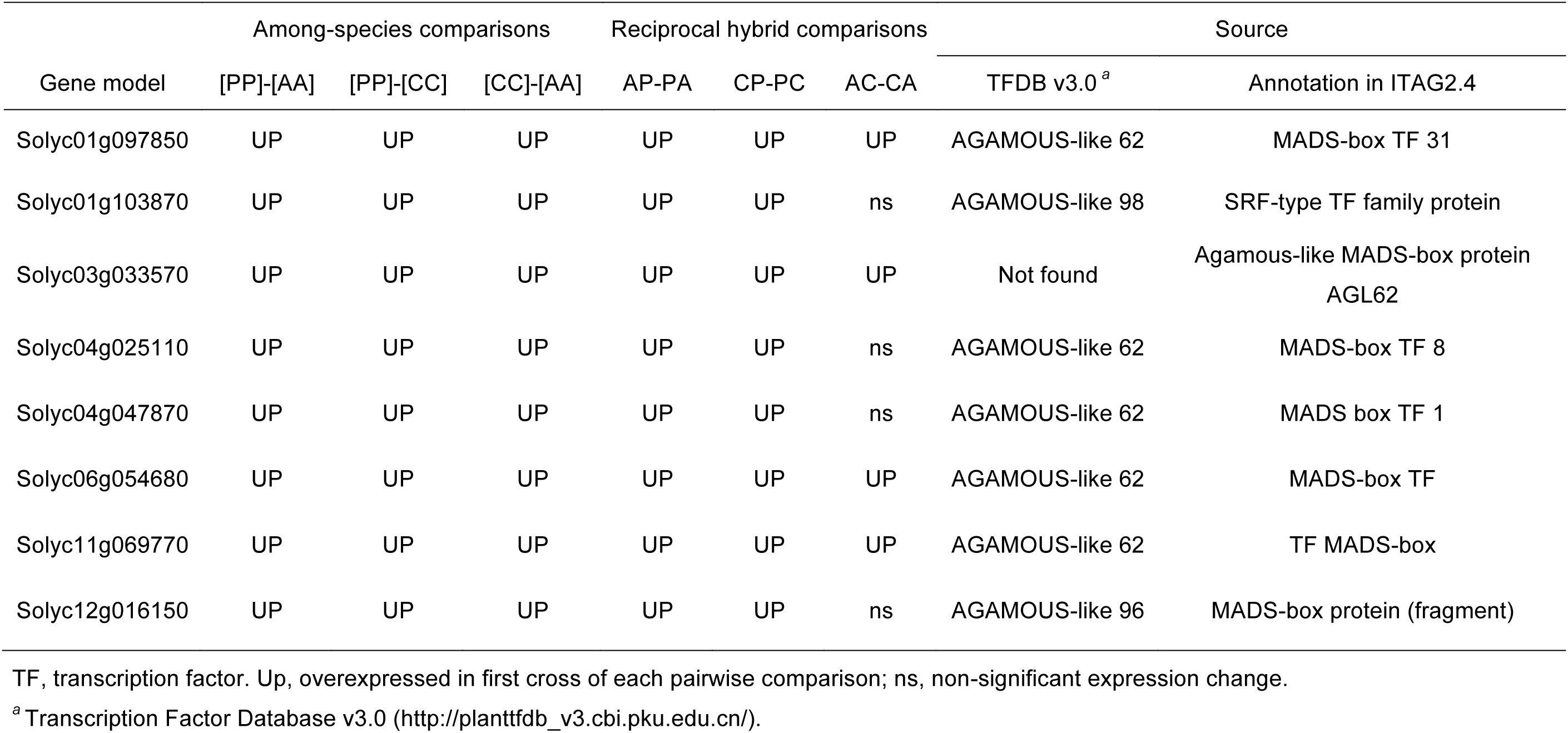
Expression pattern and annotation of eight AGL genes potentially contributing to differences in effective ploidy between species.

While genomic imprinting—which is at the core of the parental conflict theory—probably plays a crucial role in the evolution of effective ploidy (Lafon-Placette *et al*. 2018), our results indicate that the causal functional drivers might not be restricted to imprinted genes. We suggest that regulators of parent-specific expression, rather than strictly imprinted genes, might be responsible for evolutionary changes in effective ploidy. Specifically, frequent gene duplication and neofunctionalization within specific genes families such as MADS-Box TFs (*e.g.* AGL genes; Martínez-Castilla and Alvarez-Buylla 2003) and/or imprinted genes (Yoshida and Kawabe 2013; Qiu *et al.* 2014), together with epigenetic variation impacting the control of transposable elements (Pignatta *et al*. 2014; Lafon-Placette *et al*. 2018), might modify transcript abundance and expression modes over very short evolutionary timescales. This could explain why expression levels and imprinting status of specific genes vary between closely related species such as wild tomato lineages (Roth *et al*. 2018b).

It has been shown that *Arabidopsis* AGL genes act within a network and that they can be non-imprinted, maternally expressed, or paternally expressed (Walia *et al.* 2009; Bai and Settles 2015). Thus, any perturbation of expression levels among co-adapted AGLs in hybrids might be at the root of the genome-wide perturbations observed in strong-HSF hybrids. Within species, parental conflict might be stabilized by gene expression co-adaptation within functional networks, which might also determine the property ‘effective ploidy’. When parental species have accrued diverged effective ploidies this equilibrium may be disrupted in their hybrids, acting as a postzygotic reproductive barrier with varying quantitative effects depending on the disparity of effective ploidies as manifested in the endosperm. In this context, our work is the first to explore genome-wide expression correlates of dissimilar effective ploidies in the endosperm, thus enabling the exploration of possible links between parental conflict, expression dosage and HSF in flowering plants. It may also have practical applications in plant breeding, for example to enhance hybridization success between crops and their wild relatives by compensating effective ploidy differences with targeted, experimental ploidy changes.

## Acknowledgments

We are grateful to Maja Frei and Esther Zürcher for taking expert care of the plants, to Beatrice Arnold for preparing RNA-Seq libraries, and to Claudia Michel, Silvia Kobel, and Joachim Hehl for further technical help. We thank Margot Paris for her advice on experimental design and analyses and Niklaus Zemp, Stefan Zoller and Mathias Scharmann for their generous bioinformatics advice. We are grateful to Alex Widmer for his general support of this project. We thank the C.M. Rick Tomato Genetics Resource Center at U.C. Davis for generously supplying seed samples. We acknowledge the technical support on histological preparations and laser-microdissections provided by SECTION-LAB (Hiroshima, Japan) and ScopeM (ETH Zurich, Switzerland). Sequencing data were produced at the Functional Genomics Center Zurich (University of Zurich, Switzerland). We thank the Genetic Diversity Center (ETH Zurich, Switzerland) and the Swiss Institute for Bioinformatics (Lausanne, Switzerland) for providing valuable tools and training for bioinformatics analyses. This work was supported by the Swiss National Science Foundation [31003A_130702 to T.S.] and an ETH Research Grant [ETH-40 13-2 to T.S. and Alex Widmer].

## SUPPLEMENTARY INFORMATION

**Figure S1.**
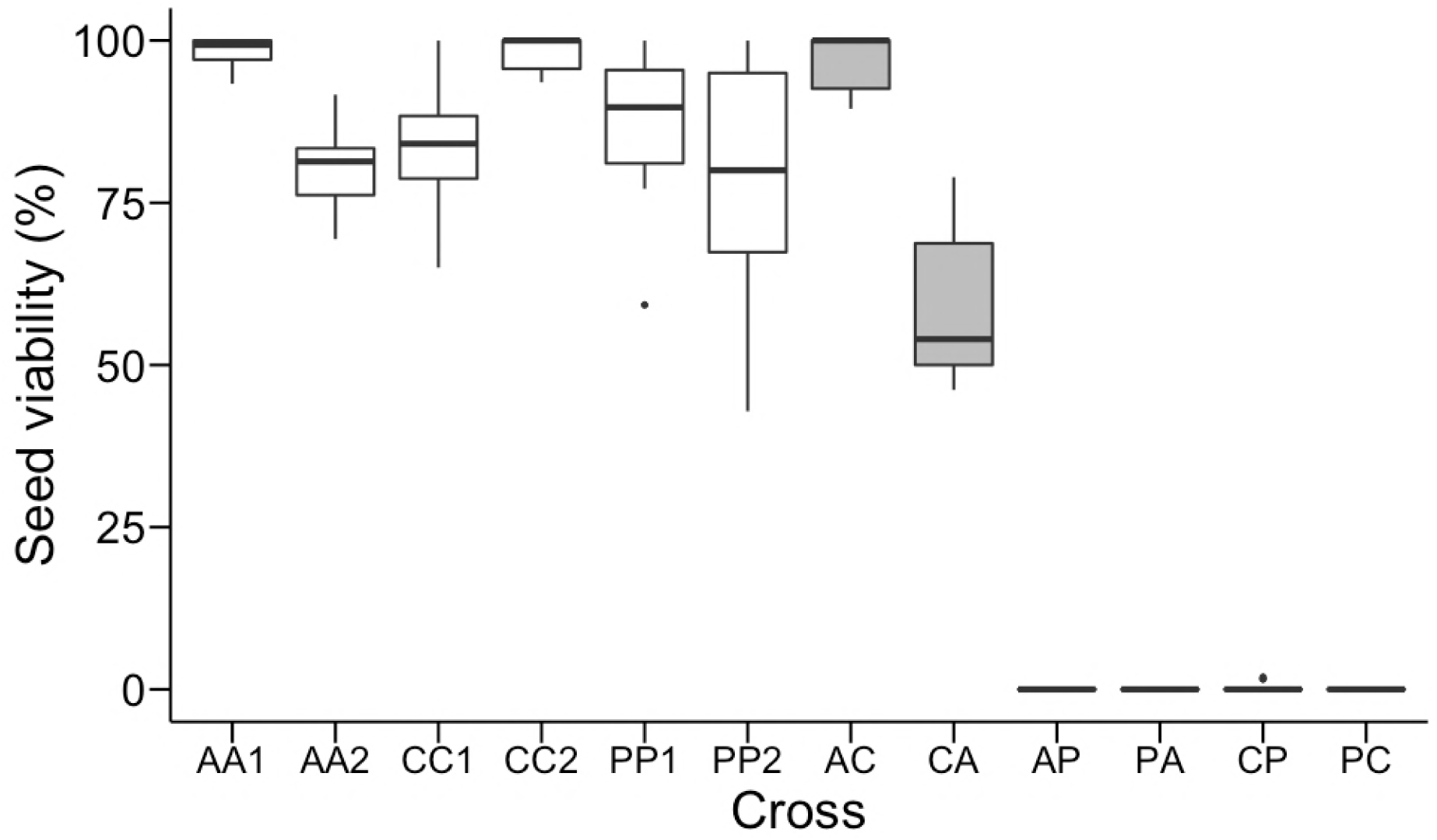
Box plot representing the distribution of seed viability in all crosses used in this study. Assessment of seed viability was performed visually at fruit maturity 60 days after pollination (data source: Roth *et al.* 2018a). Open boxes, intraspecific crosses; grey boxes, hybrid crosses. A, *S. arcanum* var. marañón; C, *S. chilense*; P, *S. peruvianum*. **AA1**, LA2185A × LA1626B; **AA2**, LA1626B × LA2185A; **CC1**, LA4329B × LA2748B; **CC2**, LA2748B × LA4329B; **PP1**, LA2744B × LA2964A; **PP2**, LA2964A × LA2744B; **AC**, LA2185A × LA4329B; **CA**, LA4329B × LA2185A; **AP**, LA2185A × LA2744B; **PA**, LA2744B × LA2185A; **CP**, LA4329B × LA2744B; **PC**, LA2744B × LA4329B. Cross specifications are identical in all other display items.

**Figure S2.**
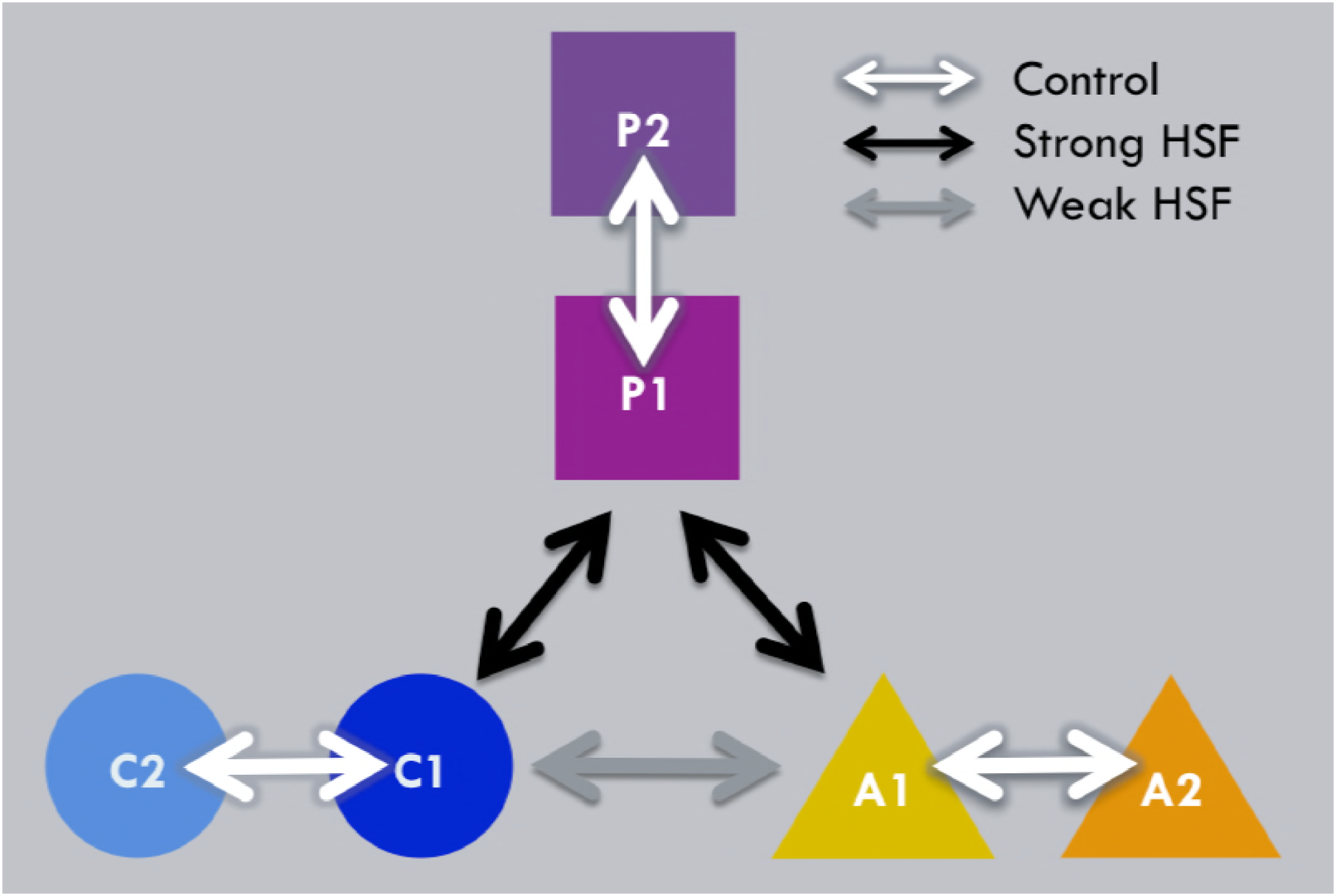
Crossing design representing the six reciprocal crosses used for our endosperm RNA-Seq experiment. Arrows represent reciprocal crosses (white, intraspecific; black, hybrid crosses resulting in strong hybrid seed failure (HSF); grey, hybrid crosses resulting in weak HSF). Each shape represents one wild tomato genotype sampled from separate populations in species A (*S. arcanum* var. marañón; triangles), C (S. chilense; circles) and P (S. peruvianum; squares). **A1**, LA2185A; **A2**, LA1626B; **C1**, LA4329B; **C2**, LA2748B; **P1**, LA2744B; **P2**, LA2964A.

**Table S1** File composed of four data sheets: ‘Contrasts’ contains the list of 18 comparisons with their corresponding contrasts used in this study; ‘DEGs’ summarizes differential gene expression (DGE) for each of them; ‘GO_enrichment’ summarizes GO-term enrichments for differentially expressed genes (DEGs) in selected categories; ‘DGE_imprinted_genes’ lists the status of candidate imprinted genes and their differential expression in all tested contrasts (separate Excel data file).

## Literature Cited

Alexa A, Rahnenführer J. 2016. topGO: enrichment analysis for gene ontology. R Package Version 2.28.0.

Anders S, Pyl PT, Huber W. 2015. HTSeq-A Python framework to work with high-throughput sequencing data. Bioinformatics 31: 166–169.

Bai F, Settles AM. 2015. Imprinting in plants as a mechanism to generate seed phenotypic diversity. Frontiers in Plant Science 5: 780.

Baroux C, Spillane C, Grossniklaus U. 2002. Evolutionary origins of the endosperm in flowering plants. Genome Biology 3: reviews1026.1.

Beddows I, Reddy A, Kloesges T, Rose LE. 2017. Population genomics in wild tomatoes—the interplay of divergence and admixture. Genome Biology and Evolution 9: 3023–3038.

Berger F, Grini PE, Schnittger A. 2006. Endosperm: an integrator of seed growth and development. Current Opinion in Plant Biology 9: 664–670.

Brandvain Y, Haig D. 2005. Divergent mating systems and parental conflict as a barrier to hybridization in flowering plants. The American Naturalist 166: 330–338.

Burkart-Waco D, Ngo K, Dilkes B, Josefsson C, Comai L. 2013. Early disruption of maternal-zygotic interaction and activation of defense-like responses in *Arabidopsis* interspecific crosses. The Plant Cell 25: 2037–2055.

Combes MC, Hueber Y, Dereeper A, Rialle S, Herrera JC, Lashermes P. 2015. Regulatory divergence between parental alleles determines gene expression patterns in hybrids. Genome Biology and Evolution 7: 1110–1121.

Cooper DC, Brink RA. 1945. Seed collapse following matings between diploid and tetraploid races of *Lycopersicon pimpinellifolium*. Genetics 30: 376–401.

Dilkes BP, Spielman M, Weizbauer R, Watson B, Burkart-Waco D, Scott RJ, Comai L. 2008. The maternally expressed WRKY transcription factor TTG2 controls lethality in interploidy crosses of *Arabidopsis*. PLoS Biology 6: 2707–2720.

Dion-Côté AM, Renaut S, Normandeau E, Bernatchez L. 2014. RNA-seq reveals transcriptomic shock involving transposable elements reactivation in hybrids of young lake whitefish species. Molecular Biology and Evolution 31: 1188–1199.

Figueiredo DD, Batista RA, Roszak PJ, Hennig L, Köhler C. 2016. Auxin production in the endosperm drives seed coat development in *Arabidopsis*. eLife 5: e20542.

Fiume E, Coen O, Xu WJ, Lepiniec L, Magnani E. 2017. Growth of the *Arabidopsis* sub-epidermal integument cell layers might require an endosperm signal. Plant Signaling and Behavior 12: e1339000.

Florez-Rueda AM. 2014. Postzygotic barriers to interbreeding in wild tomatoes: genomic imprinting and transcriptional signatures of hybrid seed failure. PhD thesis 22458, ETH Zurich, Zurich, Switzerland.

Florez-Rueda AM, Paris M, Schmidt A, Widmer A, Grossniklaus U, Städler T. 2016. Genomic imprinting in the endosperm is systematically perturbed in abortive hybrid tomato seeds. Molecular Biology and Evolution 33: 2935–2946.

Gehring M, Bubb KL, Henikoff S. 2009. Extensive demethylation of repetitive elements during seed development underlies gene imprinting. Science 324: 1447–1451.

Gehring M, Missirian V, Henikoff S. 2011. Genomic analysis of parent-of-origin allelic expression in *Arabidopsis* thaliana seeds. PLoS ONE 6: e23687.

Haig D, Westoby M. 1991. Genomic imprinting in endosperm: its effect on seed development in crosses between species, and between different ploidies of the same species, and its implications for the evolution of apomixis. Philosophical Transactions of the Royal Society B: Biological Sciences 333: 1–13.

He G, Zhu X, Elling AA, Chen L, Wang X, Guo L, Liang M, He H, Zhang H, Chen F, et al. 2010. Global epigenetic and transcriptional trends among two rice subspecies and their reciprocal hybrids. The Plant Cell 22: 17–33.

Hegarty MJ, Barker GL, Brennan AC, Edwards KJ, Abbott RJ, Hiscock SJ. 2009. Extreme changes to gene expression associated with homoploid hybrid speciation. Molecular Ecology 18: 877–889.

Hurst LD, McVean GT. 1998. Do we understand the evolution of genomic imprinting? Current Opinion in Genetics and Development 8: 701–708.

Inzé D, De Veylder L. 2006. Cell cycle regulation in plant development. Annual Review of Genetics 40: 77–105.

Jeukens J, Renaut S, St-Cyr J, Nolte AW, Bernatchez L. 2010. The transcriptomics of sympatric dwarf and normal lake whitefish (*Coregonus clupeaformis* spp., Salmonidae) divergence as revealed by next-generation sequencing. Molecular Ecology 19: 5389–5403.

John P, Zhang K, Dong C, Diederich L, Wightman F. 1993. p34^cdc2^ related proteins in control of cell cycle progression, the switch between division and differentiation in tissue development, and stimulation of division by auxin and cytokinin. Australian Journal of Plant Physiology 20: 503–526.

Johnston SA, den Nijs TPM, Peloquin SJ, Hanneman RE. 1980. The significance of genic balance to endosperm development in interspecific crosses. Theoretical and Applied Genetics 57: 5–9.

Josefsson C, Dilkes B, Comai L. 2006. Parent-dependent loss of gene silencing during interspecies hybridization. Current Biology 16: 1322–1328.

Kang I-H, Steffen JG, Portereiko MF, Lloyd A, Drews GN. 2008. The AGL62 MADS domain protein regulates cellularization during endosperm development in *Arabidopsis*. The Plant Cell 20: 635–647.

Khaitovich P, Hellmann I, Enard W, Nowick K, Leinweber M, Franz H, Weiss G, Lachmann M, Pääbo S. 2005. Parallel patterns of evolution in the genomes and transcriptomes of humans and chimpanzees. Science 309: 1850–1854.

Kinsella, Rhoda J, Kähäri A, Haider S, Zamora J, Proctor G, Spudich G, Almeida-King J, Staines D, Derwent P, Kerhornou A, Kersey P. 2011. Ensembl BioMarts: a hub for data retrieval across taxonomic space. Database 2011: bar030.

Klosinska M, Picard CL, Gehring M. 2016. Conserved imprinting associated with unique epigenetic signatures in the *Arabidopsis* genus. Nature Plants 2: 16145.

Köhler C, Hennig L, Spillane C, Pien S, Gruissem W, Grossniklaus U. 2003. The Polycomb-group protein MEDEA regulates seed development by controlling expression of the MADS-box gene PHERES1. Genes and Development 17: 1540–1553.

Köhler C, Mittelsten Scheid O, Erilova A. 2010. The impact of the triploid block on the origin and evolution of polyploid plants. Trends in Genetics 26: 142–148.

Kradolfer D, Hennig L, Köhler C. 2013. Increased maternal genome dosage bypasses the requirement of the FIS Polycomb Repressive Complex 2 in *Arabidopsis* seed development. PLoS Genetics 9: e1003163.

Labate JA, Robertson LD, Strickler SR, Mueller LA. 2014. Genetic structure of the four wild tomato species in the *Solanum peruvianum* s.1. species complex. Genome 57: 169–180.

Lafon-Placette C, Köhler C. 2016. Endosperm-based postzygotic hybridization barriers: developmental mechanisms and evolutionary drivers. Molecular Ecology 25: 2620–2629.

Lafon-Placette C, Johannessen IM, Hornslien KS, Ali MF, Bjerkan KN, Bramsiepe J, Glöckle BM, Rebernig CA, Brysting AK, Grini PE, et al. 2017. Endosperm-based hybridization barriers explain the pattern of gene flow between *Arabidopsis lyrata* and *Arabidopsis arenosa* in central Europe. Proceedings of the National Academy of Sciences U.S.A. 114: E1027–E1035.

Lafon-Placette C, Hatorangan MR, Steige KA, Cornille A, Lascoux M, Slotte T, Köhler C. 2018. Paternally expressed imprinted genes associate with hybridization barriers in *Capsella*. Nature Plants 4: 352–357.

Landry CR, Hartl DL, Ranz JM. 2007. Genome clashes in hybrids: insights from gene expression. Heredity 99: 483–493.

Leblanc O, Pointe C, Hernandez M. 2002. Cell cycle progression during endosperm development in *Zea mays* depends on parental dosage effects. The Plant Journal 32: 1057–1066.

Lester RN, Kang JH. 1998. Embryo and endosperm function and failure in *Solanum* species and hybrids. Annals of Botany 82: 445–453.

Li N, Dickinson HG. 2010. Balance between maternal and paternal alleles sets the timing of resource accumulation in the maize endosperm. Proceedings of the Royal Society B: Biological Sciences 277: 3–10.

Luo M, Taylor JM, Spriggs A, Zhang H, Wu X, Russell S, Singh M, Koltunow A. 2011. A genome-wide survey of imprinted genes in rice seeds reveals imprinting primarily occurs in the endosperm. PLoS Genetics 7: e1002125.

Martin M. 2011. Cutadapt removes adapter sequences from high-throughput sequencing reads. EMBnet J. 17: 10–12.

Martínez-Castilla LP, Alvarez-Buylla ER. 2003. Adaptive evolution in the *Arabidopsis* MADS-box gene family inferred from its complete resolved phylogeny. Proceedings of the National Academy of Sciences U.S.A. 100: 13407–13412.

Moyers BT, Rieseberg LH. 2013. Divergence in gene expression is uncoupled from divergence in coding sequence in a secondarily woody sunflower. International Journal of Plant Sciences 174: 1079–1089.

Nowack MK, Ungru A, Bjerkan KN, Grini PE, Schnittger A. 2010. Reproductive cross-talk: seed development in flowering plants. Biochemical Society Transactions 38: 604–612.

Nuzhdin SV, Wayne ML, Harmon KL, McIntyre LM. 2004. Common pattern of evolution of gene expression level and protein sequence in *Drosophila*. Molecular Biology and Evolution 21: 1308–1317.

Okonechnikov K, Conesa A, García-Alcalde F. 2016. Qualimap 2: advanced multi-sample quality control for high-throughput sequencing data. Bioinformatics 32: 292–294.

Oneal E, Willis JH, Franks RG. 2016. Disruption of endosperm development is a major cause of hybrid seed inviability between *Mimulus guttatus* and *Mimulus nudatus*. New Phytologist 210: 1107–1120.

Orozco-Arroyo G, Paolo D, Ezquer I, Colombo L. 2015. Networks controlling seed size in *Arabidopsis*. Plant Reproduction 28: 17–32.

Ortiz R, Ehlenfeldt MK. 1992. The importance of endosperm balance number in potato breeding and the evolution of tuber-bearing *Solanum* species. Euphytica 60: 105–113.

Patel RK, Jain M. 2012. NGS QC Toolkit: a toolkit for quality control of next generation sequencing data. PLoS ONE 7: e30619. doi: 10.1371/journal.pone.0030619.

Pignatta D, Erdmann RM, Scheer E, Picard CL, Bell GW, Gehring M. 2014. Natural epigenetic polymorphisms lead to intraspecific variation in *Arabidopsis* gene imprinting. eLife 3: e03198.

Qiu Y, Liu S-L, Adams KL. 2014. Frequent changes in expression profile and accelerated sequence evolution of duplicated imprinted genes in *Arabidopsis*. Genome Biology and Evolution 6: 1830–1842.

R Development Core Team. 2014. R: a language and environment for statistical computing. Vienna (Austria): R Foundation for Statistical Computing. URL: http://www.R–project.org/.

Raza MA, Yu NN, Wang D, Cao LW, Gan SS, Chen LP. 2017. Differential DNA methylation and gene expression in reciprocal hybrids between *Solanum lycopersicum* and *S. pimpinellifolium*. DNA Research 24: 597–607.

Rebernig CA, Lafon-Placette C, Hatorangan MR, Slotte T, Köhler C. 2015. Non-reciprocal interspecies hybridization barriers in the *Capsella* genus are established in the endosperm. PLoS Genetics 11: e1005295.

Renaut S, Nolte AW, Bernatchez L. 2009. Gene expression divergence and hybrid misexpression between lake whitefish species pairs (*Coregonus* spp. Salmonidae). Molecular Biology and Evolution 26: 925–936.

Renaut S, Grassa CJ, Moyers BT, Kane NC, Rieseberg LH. 2012. The population genomics of sunflowers and genomic determinants of protein evolution revealed by RNAseq. Biology 1: 575–596.

Robinson MD, McCarthy DJ, Smyth GK. 2010. edgeR: a Bioconductor package for differential expression analysis of digital gene expression data. Bioinformatics 26: 139–140.

Roth M, Florez-Rueda AM, Griesser S, Paris M, Städler T. 2018a. Incidence and developmental timing of endosperm failure in post-zygotic isolation between wild tomato lineages. Annals of Botany 121: 107–118.

Roth M, Florez-Rueda AM, Paris M, Städler T. 2018b. Wild tomato endosperm transcriptomes reveal common roles of genomic imprinting in both nuclear and cellular endosperm. The Plant Journal 95: 1084–1101.

Rushton PJ, Somssich IE, Ringler P, Shen QJ. 2010. WRKY transcription factors. Trends in Plant Science 15: 247–258.

Sabelli PA, Larkins BA. 2009. The contribution of cell cycle regulation to endosperm development. Sexual Plant Reproduction 22: 207–219.

Schruff MC, Spielman M, Tiwari S, Adams S, Fenby N, Scott RJ. 2006. The AUXIN RESPONSE FACTOR 2 gene of *Arabidopsis* links auxin signalling, cell division, and the size of seeds and other organs. Development 133: 251–261.

Scott RJ, Spielman M, Bailey J, Dickinson HG. 1998. Parent-of-origin effects on seed development in *Arabidopsis thaliana*. Development 125: 3329–3341.

Sekine D, Ohnishi T, Furuumi H, Ono A, Yamada T, Kurata N, Kinoshita T. 2013. Dissection of two major components of the post-zygotic hybridization barrier in rice endosperm. The Plant Journal 76: 792–799.

Sharma DR, Kaur R, Kumar K. 1996. Embryo rescue in plants – a review. Euphytica 89: 325–337.

Shirzadi R, Andersen ED, Bjerkan KN, Gloeckle BM, Heese M, Ungru A, Winge P, Koncz C, Aalen RB, Schnittger A, et al. 2011. Genome-wide transcript profiling of endosperm without paternal contribution identifies parent-of-origin-dependent regulation of AGAMOUS-LIKE36. PLoS Genetics 7: e1001303.

Städler T, Arunyawat U, Stephan W. 2008. Population genetics of speciation in two closely related wild tomatoes (*Solanum* section *Lycopersicon*). Genetics 178: 339–350.

Stelkens R, Seehausen O. 2009. Genetic distance between species predicts novel trait expression in their hybrids. Evolution 63: 884–897.

Stoute AI, Varenko V, King GJ, Scott RJ, Kurup S. 2012. Parental genome imbalance in *Brassica oleracea* causes asymmetric triploid block. The Plant Journal 71: 503–516.

Tellier A, Fischer I, Merino C, Xia H, Camus-Kulandaivelu L, Städler T, Stephan W. 2011. Fitness effects of derived deleterious mutations in four closely related wild tomato species with spatial structure. Heredity 107: 189–199.

The Tomato Genome Consortium. 2012. The tomato genome sequence provides insights into fleshy fruit evolution. Nature 485: 635–641.

Trapnell C, Pachter L, Salzberg SL. 2009. TopHat: discovering splice junctions with RNA-Seq. Bioinformatics 25: 1105–1111. doi: 10.1093/bioinformatics/btp120.

Valentine DH, Woodell SRJ. 1963. Studies in British primulas. X. Seed incompatibility in intraspecific and interspecific crosses at diploid and tetraploid levels. New Phytologist 62: 125–143.

von Wangenheim KH, Peterson HP. 2004. Aberrant endosperm development in interploidy crosses reveals a timer of differentiation. Developmental Biology 270: 277–289.

Walia H, Josefsson C, Dilkes B, Kirkbride R, Harada J, Comai L. 2009. Dosage-dependent deregulation of an AGAMOUS-LIKE gene cluster contributes to interspecific incompatibility. Current Biology 19: 1128–1132.

Wang L, Wang, S, Li W. 2012. RSeQC: quality control of RNA-seq experiments. Bioinformatics 28: 2184–2185.

Warnes GR, Bolker B, Bonebakker L, Gentleman R, Liaw WHA, Lumley M, Maechler M, Magnusson A, Moeller A et al. 2016. gplots: various R programming tools for plotting data. R Package Version 3.0.1.

Waters AJ, Makarevitch I, Eichten SR, Swanson-Wagner RA, Yeh C-T, Xu W, Schnable PS, Vaughn MW, Gehring M, Springer NM. 2011. Parent-of-origin effects on gene expression and DNA methylation in the maize endosperm. The Plant Cell 23: 4221–4233.

Waters AJ, Bilinski P, Eichten SR, Vaughn MW, Ross-Ibarra J, Gehring M, Springer NM. 2013. Comprehensive analysis of imprinted genes in maize reveals allelic variation for imprinting and limited conservation with other species. Proceedings of the National Academy of Sciences U.S.A. 110: 19639–19644.

Weisstein AE, Spencer HG. 2003. The evolution of genomic imprinting via variance minimization: an evolutionary genetic model. Genetics 165: 205–222.

Wilkinson L. 2011. Venneuler: Euler and Venn diagrams. R Package Version 1.1–0.

Wolf JB, Hager R. 2006. A maternal–offspring coadaptation theory for the evolution of genomic imprinting. PLoS Biology 4: e380.

Wolf JBW, Bayer T, Haubold B, Schilhabel M, Rosenstiel P, Tautz D. 2010. Nucleotide divergence vs. gene expression differentiation: comparative transcriptome sequencing in natural isolates from the carrion crow and its hybrid zone with the hooded crow. Molecular Ecology 19: 162–175.

Xu W, Fiume E, Coen O, Pechoux C, Lepiniec L, Magnani E. 2016. Endosperm and nucellus develop antagonistically in *Arabidopsis* seeds. The Plant Cell 28: 1343–1360.

Yoshida T, Kawabe A. 2013. Importance of gene duplication in the evolution of genomic imprinting revealed by molecular evolutionary analysis of the type I MADS-Box gene family in *Arabidopsis* species. PLoS ONE 8: e73588.

Zhang M, Li N, He W, Zhang H, Yang W, Liu B. 2016. Genome-wide screen of genes imprinted in sorghum endosperm, and the roles of allelic differential cytosine methylation. The Plant Journal 85: 424–436.

